# Effects of a field-sprayed antibiotic on bee foraging behavior and pollination in pear orchards

**DOI:** 10.1101/2023.02.14.528407

**Authors:** Laura Avila, Christopher McCullough, Annie Schiffer, JoMari Moreno, Neha Ganjur, Zachary Ofenloch, Tianna DuPont, Louis Nottingham, Nicole M. Gerardo, Berry J. Brosi

## Abstract

Broadcast spraying of antibiotics in crops is widely used for controlling bacterial plant pathogens. The effects of antibiotics on non-target (and especially beneficial) organisms in cropping systems, however, are not well studied. Pollinators are of particular concern because in pear and apple crops, antibiotics for controlling fire blight (*Erwinia amylovora*) are sprayed during bloom, likely exposing pollinators. This is especially relevant as laboratory evidence suggests that antibiotics could have sublethal effects on bee foraging behavior and colony health. But to our knowledge these potential impacts have not been studied in field settings. Here, we compared the effects of two fire blight control methods, a single spray of an antibiotic (oxytetracycline) and a biological antagonist (*Aureobasidium pullulans*), on honey bee (*Apis mellifera*) foraging, pollination, and fruit set in pear orchards. Complementing these field assessments, we conducted laboratory experiments to examine the effects of these treatments on locomotion and foraging behavior of the bumble bee species, *Bombus vosnesenskii*. We found that honey bees visited fewer flowers and foraged longer on each flower in orchards sprayed with antibiotics than with biological product, but there were no differences in pollination and seed set. The pear cultivars we worked in, however, can self-pollinate. In the lab, we found that feeding on high doses of either the antibiotic or the biological antagonist reduced bumble bee foraging behavior relative to controls. The limited impact of antibiotics on pear pollination observed in this study suggest that antibiotics pose a low economic risk to pear growers, especially for self-compatible cultivars. Still, crops with higher pollinator dependence may be more affected by reductions in pollinator visitation. Future studies should examine the impacts of multiple antibiotic sprays within a season, which are common during warm springs, and their long-term health impacts on both individual bees and colonies.

**Highlights:**

- Antibiotics are sprayed on many crops to control plant bacterial pathogens.
- The impacts of antibiotics on beneficial organisms in agriculture are unknown.
- We studied antibiotic impacts on bee behavior and pollination function in pears.
- Bees exposed to antibiotics visit fewer flowers and this could impact bee fitness.
- Despite decreased bee visitation, we did not detect a reduction in crop pollination.

**Graphical Abstract:** 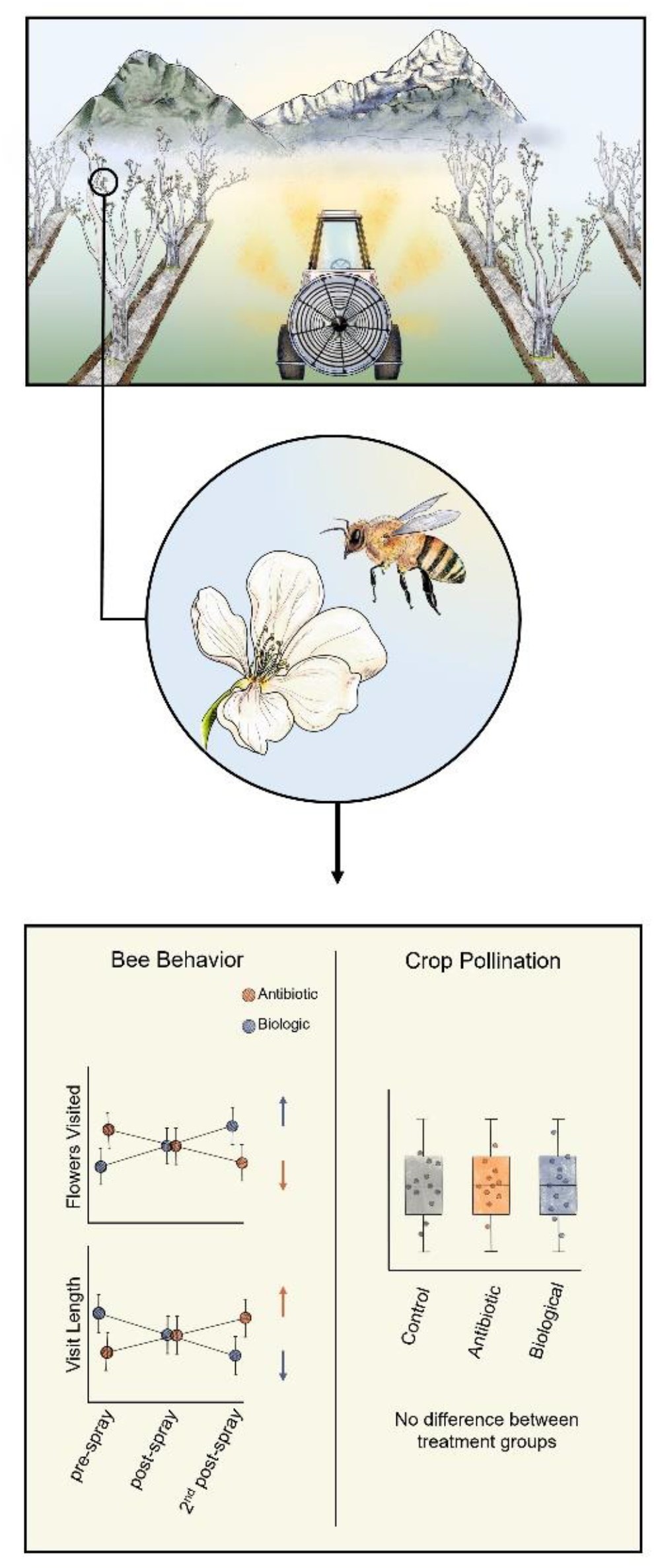

## 1. Introduction

Agricultural antibiotic use in farm animals has received a great deal of scientific and media attention (Landers et al., 2012; Manyi-Loh et al., 2018), but there is less awareness of the broadcast-spray of antibiotics in crops. Antibiotics, such as oxytetracycline, streptomycin, and kasugamycin, are a key tool in managing bacterial diseases in economically important crops globally (Taylor and Reeder, 2020). In the United States, these antibiotics are commonly used in apple (*Malus domestica*) and pear (*Pyrus communis*) orchards to prevent fire blight (*Erwinia amylovora*) (S1), a bacterial pathogen that infects trees via their flowers. Fire blight sprays occur during and near bloom, exposing pollinators and other beneficial arthropods to these chemicals. Although we are unaware of evidence of acute antibiotic toxicity to arthropods, field antibiotic exposure could have sublethal health and behavior impacts in beneficial insects. The presence of antibiotic residues may deter foraging and/or deplete insect microbial symbionts, which are known to play important roles in critical aspects of insect behavior including learning (Li et al., 2021) and locomotion (Schretter et al., 2018). Sublethal affects could have economic impacts by decreasing pollinator efficacy, leading to reduced fruit set and smaller fruit, or pest control by natural enemies. However, there have been no field-level studies examining how antibiotics may affect beneficial arthropods, particularly managed pollinators.

Several laboratory experiments have demonstrated that the consumption of two common crop antibiotics, oxytetracycline and streptomycin, could negatively impact pollinators. Studies examining these antibiotics on bees can be grouped into two categories: (1) exposure levels equal or below the maximum allowed concentrations (≤ 200 ppm), and (2) above (>200 ppm). Exposure levels in the first group (≤ 200 ppm) are not lethal to bumble bees (Marceau et al., 2021; Meeus et al., 2013), but can deplete bumble bee symbionts (Meeus et al., 2013), decrease flower visits per forager, and reduce sucrose consumption in an artificial foraging arena (Avila et al., 2022). Exposures at higher doses can decrease bumble bee (Marceau et al., 2021) and honey bee survival (Raymann et al., 2017), irreversibly deplete the honey bee symbionts (Raymann et al., 2018, 2017), impact honey bees’ ability to fight pathogens (Li et al., 2017), negatively affect honey bee olfactory learning (Zhang et al., 2020) and delay the onset of honey bee foraging (Ortiz-Alvarado et al., 2020). Beyond direct impacts on foragers consuming antibiotics, a recent study found that newly emerged honey bees can uptake the disturbed microbiome from their antibiotic-exposed nestmates (Jia et al., 2022).

Realistic field exposure levels, however, are difficult to characterize, so laboratory experiments may have used unrealistic doses. Laboratory studies do not account for factors that might influence bee exposure in the field, such as the fast degradation times of common antibiotics when exposed to heat and light (Christiano et al., 2010; Slack et al., 2021). Also, antibiotic spray concentrations can vary considerably due to differences in water volume used for pesticide applications. For example, in tree fruit, water volumes for airblast sprays commonly vary from 935.4 or 1870.8 liters per hectare based on tree size, meaning the concentration of a pesticide in spray solution can increase or decrease by half depending on the orchard (Washington State University, 2023). This concern is salient because, on the one hand, exposure levels could be less than dosing levels, particularly if there is antibiotic degradation in the field; on the other hand, exposure could be effectively greater if there is bioaccumulation of antibiotics or concentration of antibiotics (e.g., evaporation of exposed nectar). Field experiments are thus essential to understand the implications of antibiotic exposure for bee behavior and crop pollination.

Fire blight treatment in pome (apple and pear) orchards is an important system to investigate whether field-sprayed antibiotics impact bee foraging behavior and concomitant pollination services. Fire blight is a devastating bacterial disease that enters the vascular system of the fruit tree through floral nectaries (Johnson and Stockwell, 1998). Infected trees will experience death of individual twigs, limbs, or entire trees. Conventional management of fire blight relies on sprays of antibiotics (streptomycin, oxytetracycline, and kasugamycin), especially during warm and wet springs (DuPont et al., 2019). These compounds are applied over entire trees via airblast sprayers during bloom to prevent the pathogen from infecting flowers. These applications happen at a time when pome fruit growers bring honey bee colonies to pollinate their crops (Lawrence, 2022, Figure S2). Consequently, foraging bees are likely exposed to antibiotics topically (exposure during a spray application) or via consumption of antibiotic residues in floral pollen and nectar. Both foraging and non-foraging bees (including bee larvae) could be exposed to antibiotics through secondary products of nectar and pollen, such as bee bread and honey (Al-Waili et al., 2012).

An additional facet of fire blight management that makes it particularly amenable to study its impacts on pollinators is that there are non-antibiotic alternatives (Kurtulus Bastas et al., 2022), which allow for field contrasts with antibiotic use. In particular, the antagonist yeast *Aureobasidium pullulans* (Johnson et al., 2022; Ngugi et al., 2011; Sundin et al., 2009)—which has been adopted in commercial orchards with both conventional and organic management—is one common biological control method. While the impacts of *A. pullulans* on pollinator survival and behavior in the field are unknown, greenhouse studies have shown that it does not impact foraging behavior in bumble bees (Iqbal et al., 2022) nor cause lethal effects on honey bees (European Food Safety Authority, 2013). Such alternative provides a control to which antibiotic impacts can be compared.

In this study, we assessed various indicators of pollination and pollinator health in pear orchards sprayed with either antibiotics or biologicals. We measured 1) field honey bee foraging, 2) pollination, and 3) agronomic outcomes (e.g., fruit and seed set). We also measured the agronomic outcomes in three additional unsprayed, conventional orchards. This was possible as the risk of fire blight was low during the cold spring of 2022, when we conducted field work. Additionally, we assessed the impact of field exposure on 4) bumble bee locomotion (time in movement) and foraging (number of approaches to reward) in the lab relative to controls. We hypothesized that field antibiotic exposure would have negative impacts in bee behavior as shown in laboratory studies. We thus predicted that honey bees exposed to antibiotics in pear orchards would visit fewer flowers compared to those exposed to the commercial yeast antagonist. We focused on floral visitation, as it is typically used as a measure of sublethal impacts of agrochemicals on the foraging activity of bees (Stanley and Raine, 2016; Tamburini et al., 2021). Given the need for conspecific pollen transfer for the pollination of pear trees (Webster, 2002), we hypothesized that a decline in floral visitation would lead to reduced pollination and lower fruit production. In terms of bumble bee foraging outcomes in the lab, we hypothesized that antibiotic exposure would negatively impact foraging, in parallel with our hypothesis for honey bees (Avila et al., 2022; Ortiz-Alvarado et al., 2020). We thus predicted that antibiotic-fed bumble bees would move and forage less compared to the biological and control treatment.

## 2. Materials and Methods

### 2.1 Experimental sites

We used 13 commercial pear orchards within <50 km of Wenatchee, Washington State, United States, to assess the potential impacts of antibiotic sprays on pollinator behavior and pollination success (see table S22 with sampling details for each site). Due to the nature of working in commercial orchards, sites varied in general architecture, tree ages, cultivars ratios, and management practices. For our experiments, the independent variables separating orchards into treatment categories were fire blight spray materials: five conventional orchards were sprayed once with an antibiotic (‘Mycoshield’, 200 ppm oxytetracycline, applied at a rate of 1.121 kg of commercial product containing 17% oxytetracycline in 934.6 L of water per hectare), five were organic orchards sprayed once with a biological antagonist (‘Blossom Protect’, *Aureobasidium pullulans* strain DSM 14940 and DSM 14941, applied at a rate of 11.5 kg of commercial product together with 10.5 kg of Buffer Protect in 1000 L water per ha). While there could be confounding effects of comparing organic to conventional orchards (especially through the effects of other agrochemicals), there are several reasons why we believe this to be of low consequence to our experiments. First, managed bees are placed in orchards only during bloom, at which point growers stop using insecticides to avoid harming to these managed pollinators. Therefore, managed pollinators in our experiments would not be affected, directly, by variability among conventional and organic spray programs. Additionally, we monitored honey bee foraging behavior before and after fire blight sprays to reduce the shortcomings of lacking control (unsprayed) sites. While our contrast is admittedly imperfect, it represents an important first step in understanding field-realistic impacts of antibiotic exposure on pollinator behavior, pollination, and crop production. For the agronomic assessment, we included three additional conventional orchards untreated against fire blight. Hereafter, treatments will be referred to as “antibiotic”, “biological”, or “untreated control”, respectively. Most of the orchards were located at least a mile apart, though two sets of two orchards were a half mile away from one another and separated by a major road and a major river, respectively. These landscape features should minimize overlapping foraging ranges across sites, as honey bees are known to fly along the length of these landmarks if encountered during foraging flights (Collett and Graham, 2015; Menzel et al., 2019). Orchards were 5 to 25 acres in size and each orchard had a mix of Bartlett, Bosc, and D’Anjou (Green and Red) pear cultivars.

### 2.2 Pollinator foraging behavior: field

In each of our fields, growers used rented honey bee colonies for pollination. We coordinated sampling times with growers, based in part on the timing of fire blight treatment applications. All behavioral measurements took place from April 20, 2022 to May 4, 2022, during peak bloom when pollinator activity was high and before the application of floral thinning products. We assessed field honey bee foraging behavior within 48 h before and after fire blight sprays, hereafter “pre” and “post”. Measurements were conducted by walking an arbitrary path through orchards and visually documenting the foraging behavior of 25 honey bees. For each honey bee observed, we documented the number of flower landings and flower departures, using a voice recorder. We stopped following the focal bee when we lost track of it or when it flew beyond eyesight (minimum observation duration: 20 seconds; maximum: 229 seconds). Data were transcribed from recordings in the lab and used to estimate the visit length per flower per bee, and the total number of flowers visited per bee. Due to lack of honey bee activity, we could not monitor pre-spray foraging activity at two biological sites and at one antibiotic site. We monitored a second time post-spray (96 to 168 h later) at three orchards in each treatment, hereafter “2^nd^ post”. Due to weather, we were able to record the foraging activity of only two bees at one of the antibiotic-sites, and therefore excluded those data from the analysis. Due to the small number of site replicates per fire blight treatment, we analyzed the data by individual foraging bout. In total, the models included observations of 443 honey bees.

During each sampling visit, we also performed timed honey bee counts on flowers. First, we estimated mean flower density by counting the number of open flowers in a cubic meter of branches on a single tree. We then estimated honey bee abundance by counting all new bees entering the one cubic meter area over a 10-minute period; only honey bee abundance was analyzed due to low abundance of other taxa. Three counts, each on a different tree, were performed per site per sampling visit.

### 2.3 Pollination outcomes

During successful fertilization, the ovary and receptacle of the flower enlarges (Figure S3), and seeds develop (Nishitani et al., 2012). In many plant species, fertilization is dependent on transfer of outcross pollen by pollinating insects. The pear cultivars at our sites, however, exhibited a gamut of dependence on insects for fertilization and fruit development. To account for self-pollination and parthenocarpy, we bagged tree branches with flowers and later compared pollination success in branches that had stayed bagged vs. those that had bags removed (i.e., open to pollinator activity). Specifically, once flowers had reached green cluster to white bud stage, we randomly selected 40 trees per site in the center rows of each orchard. On each tree, we tagged a branch that we left “open” (i.e., did not bag). Additionally, on 20 of these trees, we bagged a branch with a muslin bag (30.5 cm × 40.6 cm) to prevent insect-mediated cross-pollination. Our experimental sites had a mix of cross-pollinated cultivars such as D’Anjou pears and self-pollinated and/or parthenocarpic cultivars such as Bartletts (Burts and Kelly, 1960; Claessen et al., 2019). In total we tagged at least 60 branches per site (∼40 open branches and ∼20 bagged branches). All the selected branches were ∼1.2 to 1.5 m high. We counted the number of clusters per branch, as clusters can be predictive of fruit pollination (Colda et al., 2021), and the number of flowers per cluster. Then, pollination success was compared among treatments using two metrics: number of pollen grains deposited per stigma and proportion of flowers with ovule expansion, herein ‘pollen deposition’ and ‘ovule expansion’, respectively. To measure pollen deposition, 24-36 hours after fire blight applications, we removed bags from 10 of the “bagged” branches per site; we refer to these as “unbagged” branches. Then, 72 to 96 hours after bag removal, we collected one open flower from each of two clusters of unbagged branches and removed and mounted the floral stigmas on slides with fuchsin jelly (Kearns and Inouye, 1993). To quantify pollen deposition, we returned flowers to the lab and counted the number of stigmas mounted and pollen grains deposited on each by scanning the entire slide under a compound microscope at 100x magnification. We analyzed a total of 120 flowers. Finally, we measured ovule expansion at petal fall by visually inspecting and counting the number of flowers with visibly enlarged ovules (an indicator of successful pollination and the first stage in fruit development) on all open, bagged and unbagged branches (Figure S3). We calculated proportion ovule expansion as the number of flowers with ovule expansion divided by the number of flowers initially counted in that branch.

### 2.4 Agronomic outcomes associated with pollination

We harvested fruits from either Green or Red D’Anjou cultivars, because these are thought to be the most pollen-limited of the varieties in our study orchards. There were no differences in terms of width and size between the red and green varieties. Once pears had reached full development (from September 13, 2022 to September 27, 2022), we collected five fruits from each of ten trees per site (a total of 50 fruits for each of the 13 sites); both fruits and trees were arbitrarily selected and harvested from branches at ∼1.2 to 1.5 m height. We stored the fruits in a cold room until processing. Fruit quality, specifically weight, size, and length, were assessed in a packing line at the Wenatchee Tree Fruit Research and Extension Center (WTFREC) of Washington State University. Afterwards, fruits were split on their equatorial axis to count mature seeds (seed set) and hand-measure fruit diameter.

### 2.5 Pollinator foraging behavior: lab

To further assess impact of the fire blight treatments on foraging behavior, we placed *Bombus vosnesenski* (Koppert, USA, Michigan) colonies in each experimental site with the intent to assess behavior in the field, however, it was extremely rare to see bumble bees visiting pear blossoms, which is consistent with other reports (e.g., van den Eijnde, 1996), and thus we were unable to assess an impact on bumble bee foraging in the field. Therefore, we carried out laboratory experiments to assess treatment effects on *B. vosnesenski* foraging behavior. We set up a completely randomized block design with *B. vosnesenskii* colonies in a greenhouse at the WTFREC. We had four quads (blocking factor) with four colonies each. Three quads had been in unsprayed control sites and one at a biological site. To equalize their management history, we brought them into the greenhouse and fed them a diet of 1 Mol sucrose solution and bee pollen (Koppert, USA, Michigan) for a month prior to the experiment. This time allowed new bees to emerge under an equivalent diet and greenhouse environment. After a month, each colony within a quad was randomly given one of four ad-libitum diets for three days: the same sucrose diet (control), sucrose plus full dose (200 ppm) of oxytetracycline, sucrose plus half dose of the antibiotic, or sucrose plus full rate of the biological product sprayed in the field. After three days, five naïve bees (i.e., bees that had never foraged outside) per colony (*n* = 20 bees per treatment) were placed in artificial foraging arenas consisting of a 10 mm glass Petri dish with a dot of honey in the middle (Figure S4). Colonies were cooled for about an hour beforehand and the bees were given 15 minutes to warm up before the assay started. The assays were conducted over two days with two quads assayed per day. We used an EthoVision XT System (Noldus Information Technology, Leesburg, VA, USA) to record videos of each bee for 45 minutes. We processed the videos with the EthoVision software and estimated metrics related to locomotion (time in movement) and foraging (number of approaches to reward). A similar set up has been used by others to assess the impact of pesticides on bees (Ingram et al., 2015; Teeters et al., 2012).

### 2.6 Data analysis

We analyzed the data in R v. 3.6.0 (R Development Core Team). We used Generalized Linear Mixed Effect Models (GLMMs) implemented with the ‘glmmTMB’ function from the “glmmTMB” package (Brooks et al., 2017). GLMMs were chosen because all models included repeated measures, with multiple non-independent measurements within each site (sometimes multiple measurements on a single tree) for field data, and different quad origins in the bumble bee laboratory study. We treated all random effects as random intercepts. We built models to assess the main effect of fire blight treatment on 1) field honey bee foraging (number of flowers visited and associated visit length); 2) pollination (pollen deposition and ovule expansion); 3) agronomic variables (fruit quality and seed set); and 4) lab bumble bee locomotion (time in movement) and foraging (approaches to reward).

We assessed two components of honey bee foraging: number of flowers visited and visit length. We modeled flowers visited (count variable) with negative binomial errors distribution. Fixed effects included interactions between fire blight treatment and sampling time (pre, post and second post), mean honey bee abundance and mean flower density. We included an offset corresponding to the length of the honey bee observation, and site as random effect (random intercept). We modeled the log of visit length (continuous variable) with Gaussian errors. We included the same fixed effect as for the number of flowers visited but removed mean honey bee abundance to decrease collinearity (variance inflation factor <5) and added site as random effect.

We also assessed two components of pollination: stigmatic pollen deposition and ovule expansion. We modeled stigmatic pollen deposition (count variable) with negative binomial errors. We included fire blight treatment as the sole fixed effect, and unique tree number nested within site as random effects. We modeled ovule expansion as a proportion, using a beta distribution to account for non-normal residual. We initially tried to model this outcome with a binomial distribution, however, the model residuals were not normally distributed. Therefore, we used a beta distribution instead. As beta models do not accept proportions equal to 1 (e.g., perfect pollination), we replaced 1 values with 0.999. Fixed effects were fire blight treatment, branch treatment (bagged, unbagged or open branch), flower clusters per branch and unique tree number nested within site as random effect.

For the agronomic outcomes, we modeled the number of mature seeds with negative binomial errors, and fruit diameter and weight with gaussian errors. For all of the agronomic outcomes, we specified fire blight treatment and cultivar (Red or Green D’Anjou) as fixed effects and site as a random effect. We log and square-transformed diameter and weight, respectively, to improve model residuals.

For the lab bumble bee outcomes, we modeled foraging (number of approaches to the reward) with negative binomial errors and locomotion with Gaussian errors. For both, we used fire blight treatment as the sole fixed effect, and unique colony number nested within quad as random effects.

We checked all models for heteroscedasticity of residuals with the ‘simulateResiduals’ function from the model “DHARMa” package (Hartig, 2020) and collinearity with the ‘check_collinearity’ function from the “performance” package (Lüdecke, 2020). We interpreted the statistical significance of fixed effects with type II or type III errors via Wald *X*^2^ tests implemented with the ‘ANOVA’ function from the “car” package (Fox et al., 2022). Whenever possible, we estimated marginal and conditional R^2^ via the ‘r.squaredGLMM’ function from the “MuMIN” package (Barton, 2022).

## 3. Results

### 3.1 Pollinator foraging behavior: field

Honey bee foraging trends were affected by fire blight treatments. According to our model predictions, the number of flowers visited per bee decreased after the antibiotic spray (0.69 less visits during observation period), while the opposite was true after the biological spray (Figure 1A; interaction Wald *X*^*2*^_(2, N = 443)_ = 16.80, *p* < 0.0002; see table S1 for full GLMM results & S2 for Wald *X*^*2*^_(2, N = 443)_ results). Honey bee visit length per flower increased after antibiotic applications and decreased after the biological spray (Figure 1B; interaction Wald *X*^*2*^ =8.911, *p* = 0.01161, see table S3 for full GLMM results & S4 for Wald *X*^*2*^ results).

**Figure 1.**
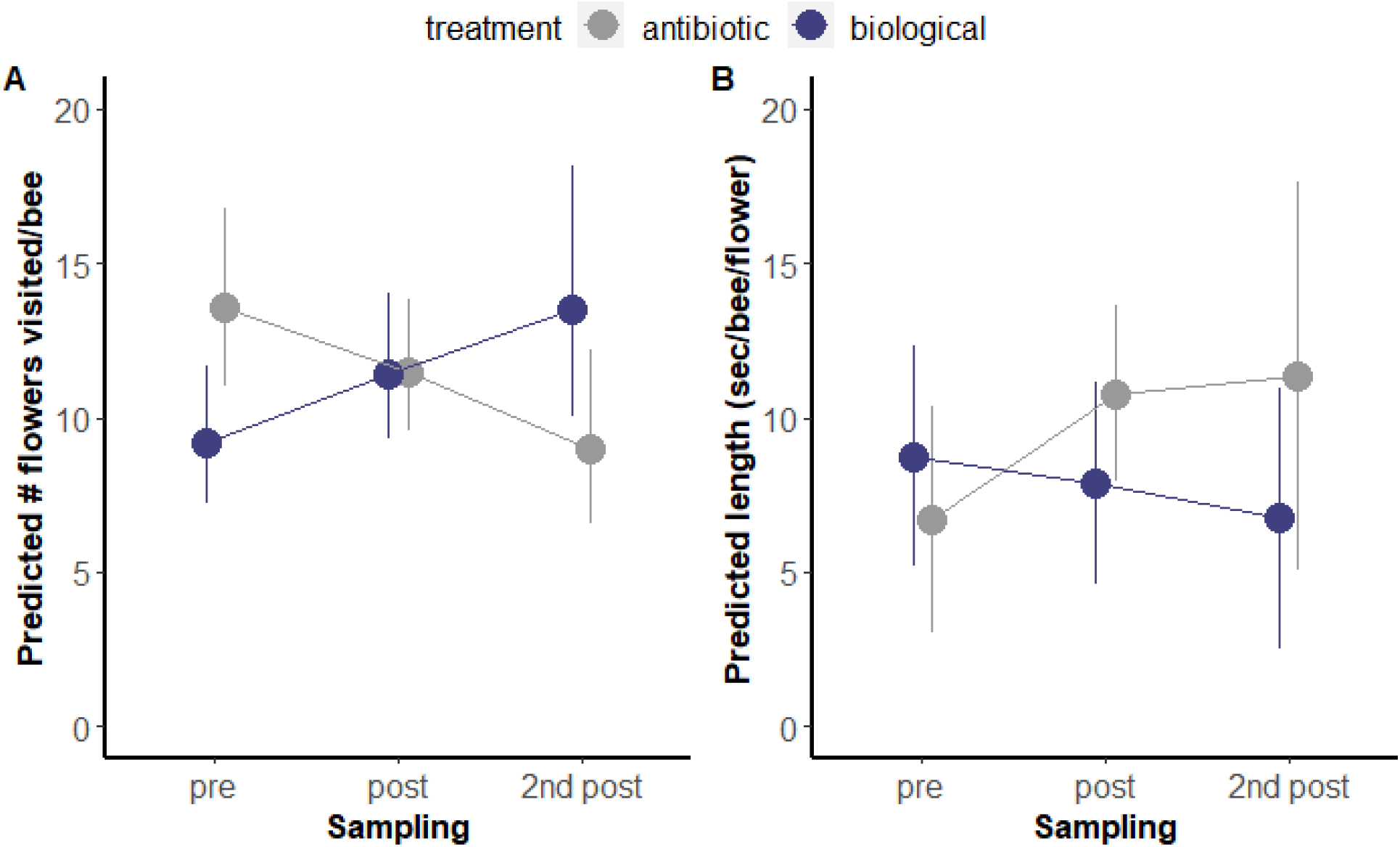
Antibiotic exposure led to a decrease in the number of flowers visited; the opposite was true for the biological product. Circles indicate means plus 95% CI. ‘Pre’ refers to 48 hrs before spray, ‘post’ to 48 hrs after spray, and ‘2^nd^ post’ to 96 to 168 hrs after spray. **A)** Predicted number of flowers visited by each honey bee forager (GLMM, interaction Wald *X*^*2*^ _(2, N = 443)_= 16.80, *p* < 0.0002); **B)** Predicted visit length (GLMM, interaction Wald *X*^*2*^_(1, N = 443)_ =8.911, *p* = 0.01161).

### 3.2 Pollination outcomes

Pollination outcomes were similar for antibiotic-sprayed and biological-sprayed sites. There were no differences in pollen deposition 24 to 36 hours after fire blight treatments (Figure 2A; GLMM, Wald *X*^*2*^_(1, N =117)_ = 0.94, *p* = 0.8490; see Table S5 for full GLMM results & S6 for Wald *X*^*2*^ results). Similarly, there was no difference on ovule expansion among treatments (Figure 2B; GLMM, Wald *X*^*2*^ _(1, N = 510)_=0.86, *p* = 0.3539; see table S7 for full GLMM results & S8 for Wald *X*^*2*^ results). Bagged branches had significantly fewer fertilized flowers than open branches (IR=1.94; CI=1.43-2.62; *p* < 0.001). Parthenocarpy and self-pollination, accounted for close to 40% of pollination across all orchards and treatments (36.6±26.4% for bagged branches vs 60.3±25.8% for open branches as shown in table S21).

**Figure 2.**
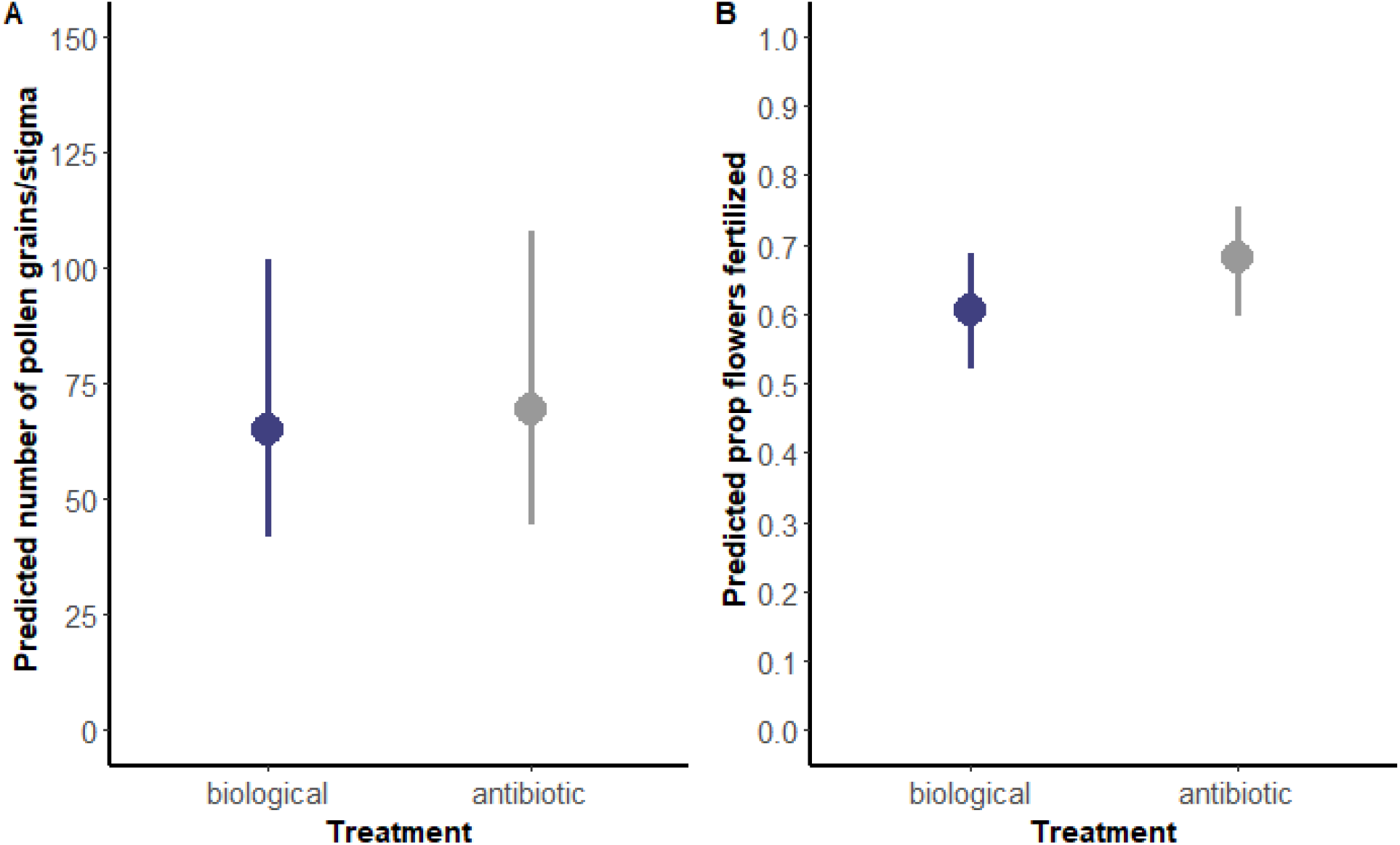
Antibiotic exposure did not impact pollen deposition or ovule expansion. For both panels, data from unbagged branches exposed to bee foraging 24-36 hours after fire blight treatment applications. **A)** Predicted number of pollen grains deposited per stigma (GLMM, Wald *X*^*2*^_(1, N =117)_ = 0.94, *p* = 0.8490); **B)** Predicted proportion of flowers with ovule expansion (GLMM, Wald *X*^*2*^_(1, N = 510)_ =0.86, *p* = 0.3539).

### 3.3 Agronomic outcomes associated with pollination

Agronomic outcomes were unaffected by fire blight treatment. There were no differences in the number of mature seeds from flowers among fire blight treated sites (Fig 3A, GLMM, Wald *X*^*2*^ _(2, N =648)_= 0.059, *p* = 0.9711). The cultivar (Green vs. Red D’Anjou) did not have a statistically significant effect on the number of mature seeds per fruit (GLMM, Wald *X*^*2*^_(1, N =648)_ = 0.874, *p* = 0.3499; see table S9 for full GLMM results & S10 for Wald *X*^*2*^ results). Fruit diameter was not affected by treatment (Fig 3B, Wald *X*^*2*^ _(2, N =648)_ = 0.347, *p* = 0.8407) or cultivar (GLMM, Wald *X*^*2*^_(1, N =648)_ = 0.336, *p* = 0.5622). Number of seeds had a statistically significant effect on fruit diameter; the more seeds the greater the diameter (Wald *X*^*2*^_(1, N =648)_ = 0.19277, *p* < 0.0001; see Figure S6 and table S12 for full GLMM results). Last, fruit length and weight were not affected by treatment (Figure 4A, Wald *X*^*2*^_(2, N =648)_ = 2.39, *p* = 0.30144 & Figure 4B, Wald *X*^*2*^ _(2, N =648)_ = 0.383, *p* = 0.825550) or cultivar (Wald *X*^*2*^_(1, N =648)_ = 1.060, *p* = 0.303322 & GLMM & *X*^*2*^_(2, N =648)_ = 0.201, *p* = 0.654162, see table S13 & S15 for full GLMM results and S14 & S16 for Wald *X*^*2*^ results).

**Figure 3.**
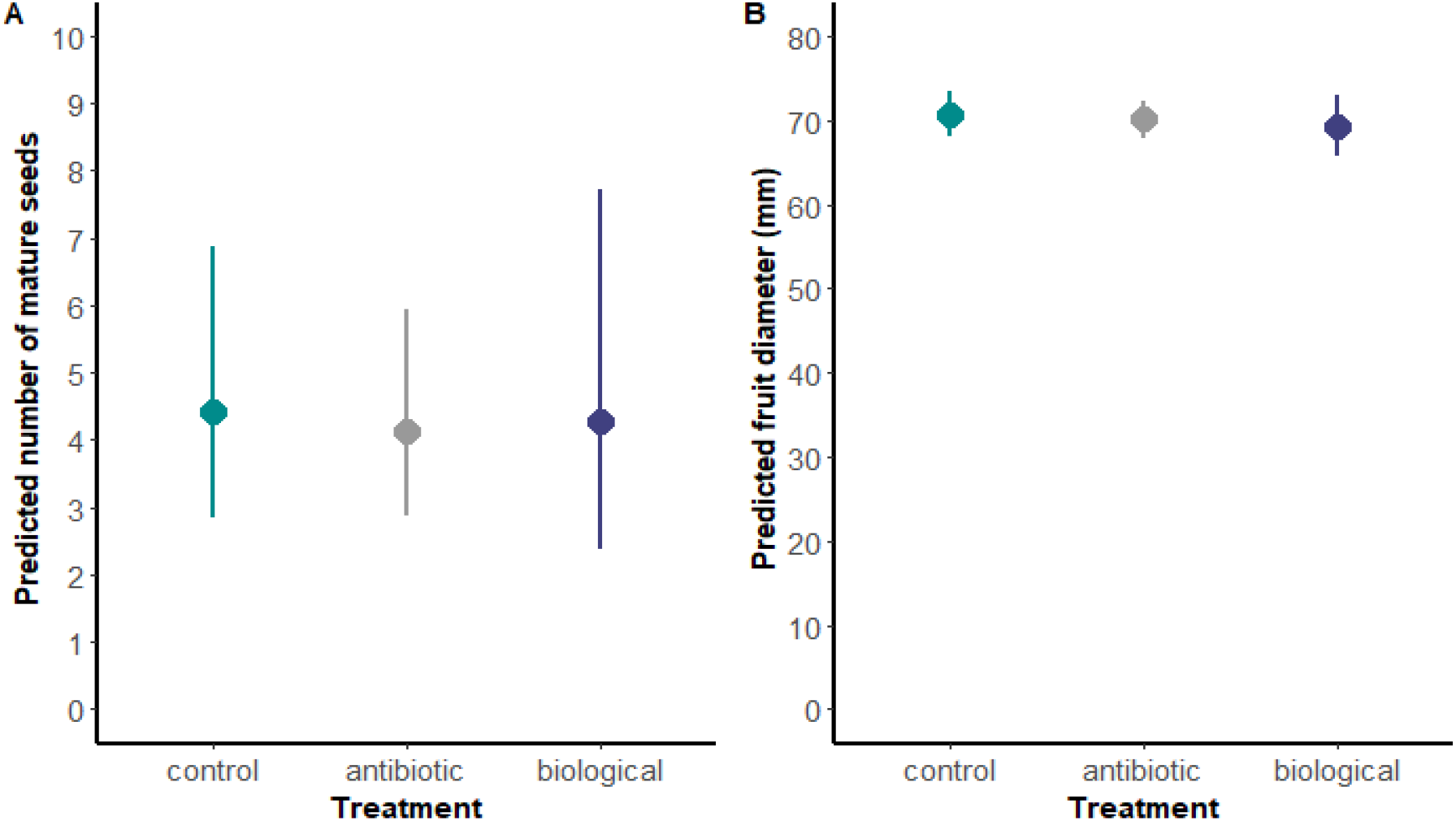
Fire blight treatments did not impact fruit production. **A)** Predicted number of mature seeds per fruit (GLMM, Wald *X*^*2*^_(2, N =648)_ = 0.059, *p* = 0.9711); **B)** Predicted diameter of fruits (GLMM, Wald *X*^*2*^_(2, N =648)_ = 0.347, *p* = 0.8407).

**Figure 4.**
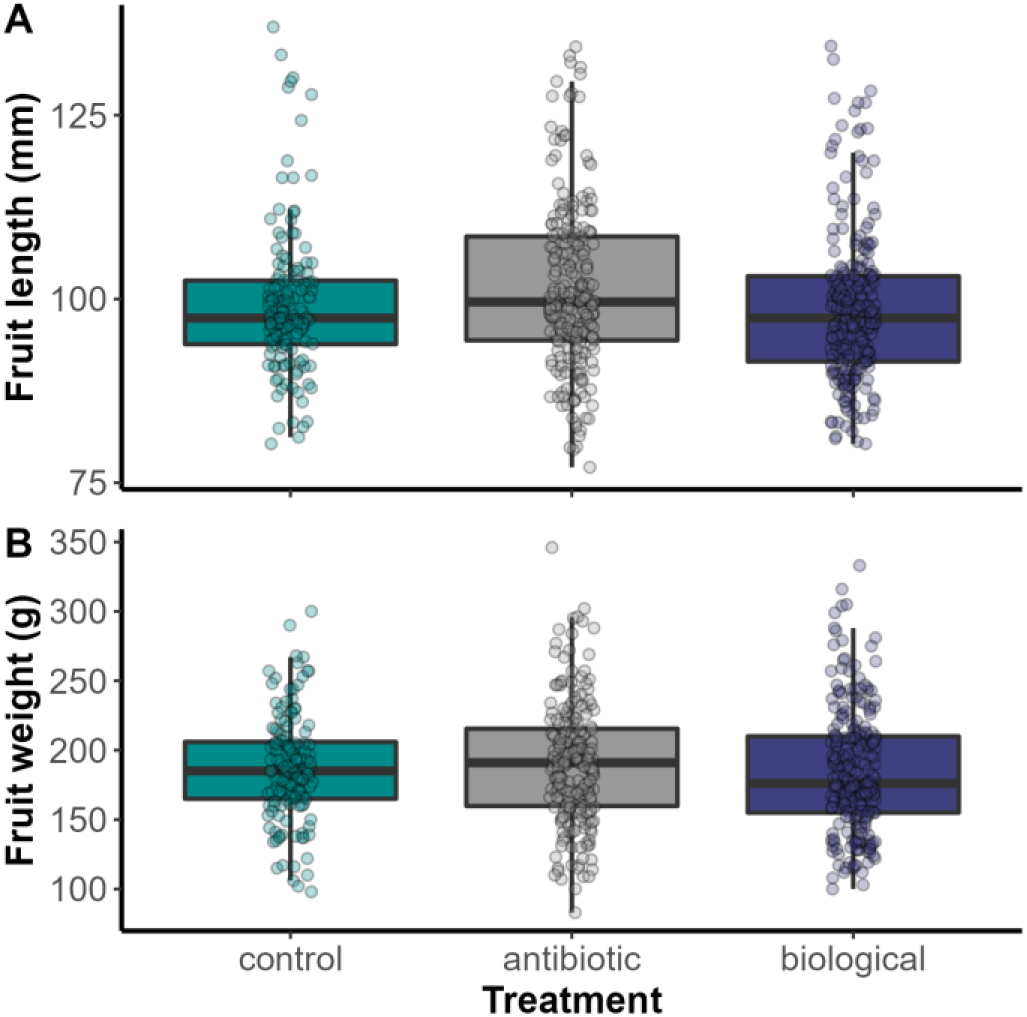
Fire blight treatments did not impact pear fruit quality. Boxplots showing interquartile ranges. Dots represent individual fruits. **A)** Measured fruit length (Figure 4A, GLMM, Wald *X*^*2*^_(2, N =648)_ = 2.39, *p* = 0.30144); **B)** Measured fruit weight (GLMM, Wald *X*^*2*^_(2, N =648)_ = 0.383, *p* = 0.8255).

### 3.4 Pollinator foraging behavior: lab

Fire blight treatments impacted bumble bee reward-seeking behavior in lab experiments. Treatment had a significant effect on the number of bumble bee foraging. Bees from both antibiotic and biological treatment groups made fewer approaches to the honey reward than those fed only water (Figure 5A; Wald *X*^*2*^ _(3, N = 80)_= 10.28, *p* = 0.01636; see table S17 for full GLMM results & S18 for Wald *X*^*2*^ results). There was no significant difference between treatments in the amount of time bees spent moving within the experimental arena (Wald *X*^*2*^ _(3, N = 80)_=3.91, *p* = 0.27180, see table S19 for full GLMM results & S20 for Wald *X*^*2*^ results). Although it was not significant, bees fed the biological fire blight material in the lab were more active than those fed the antibiotic diets (Fig 5B), which was also observed in our field experiments.

**Figure 5.**
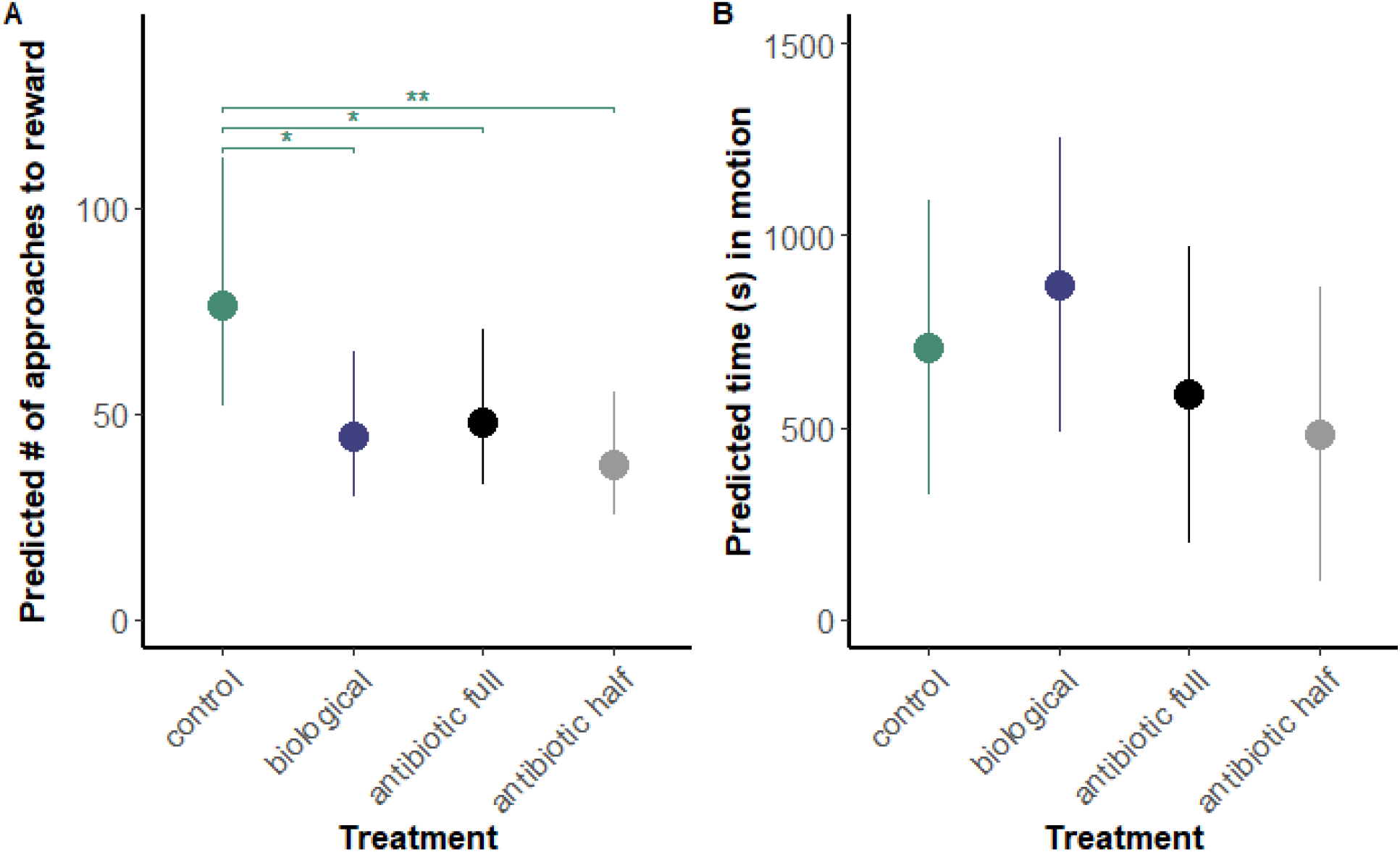
Bumble bee reward-seeking behavior changes when fed fire blight treatments. Bees were video recorded for 45 minutes inside artificial foraging arenas. Circles indicate means plus 95% CI. Asterisks indicate statistically significant differences against control, with * = *p* < 0.05; ** = *p* < 0.01. **A)** Predicted number of approaches to a honey reward by *Bombus vosnesenskii* foragers (GLMM, Wald *X*^*2*^_(3, N = 80)_ = 10.28, *p* = 0.01636); **B)** Predicted number of seconds in motion (GLMM, Wald *X*^*2*^_(3, N = 80)_ = 3.91, *p* = 0.27180).

## 4. Discussion

We found that field antibiotic exposure impacted pollinator visitation to crop flowers, but we did not find any detectable effects on pear pollination. Specifically, we found that honey bees visited fewer flowers in orchards sprayed with an antibiotic fire blight material than those sprayed with a biological fire blight material (Fig 1A). Additionally, lab bumble bees (*B. vosnesenskii*) exposed to fire blight to both antibiotic and biological treatments were less likely to forage than untreated bees (Fig 5A). Despite indications of adverse impacts on bee behavior, we found no impact of treatments on pear pollination directly, including stigmatic pollen deposition (Fig 2A) and ovule expansion (Fig 2B), or indirectly to agronomic variables such as mature seed number (Fig 3A), fruit diameter (Fig 3B), fruit length (Fig 4A), or fruit weight (Fig 4B). Many pear cultivars, however, can self-pollinate, reducing the likelihood of detecting changes to fruit production. Moreover, the levels of antibiotic exposure in our study were lower than typical (a single antibiotic spray in each orchard), and the cold weather at bloom time likely also impacted both bee foraging behavior and aspects of pear pollination, fertilization, and fruit development.

Various mechanisms may have led to the results observed in honey bee field foraging and bumble bee behavior in the lab. First, exposure to antibiotics in floral rewards could have depleted bee gut symbionts, leading to adverse changes in honey bee foraging behavior. Studies have shown that gut microbes can influence bee learning, memory (Cabirol and Haase, 2019; Li et al., 2021) and social interactions (Liberti et al., 2022), that tetracycline antibiotics used to treat foulbrood can disrupt the honey bee gut microbial community (Jia et al., 2022; Raymann et al., 2017; Soares et al., 2021), and that tetracyclines can also lead to delayed foraging in honey bee colonies (Ortiz-Alvarado et al., 2020). A very recent study showed that tetracycline exposure during adulthood delayed the expression of genes involved in the onset of foraging (Ortiz-Alvarado et al., 2022). Second, direct antibiotic intoxication could explain differences. For example, antibiotic intoxication can lead to motor deficits in vertebrates (Grill and Maganti, 2011), but this topic has not been directly studied in invertebrates to our knowledge. Third, antibiotic sprays on floral structures could have deterred bees from foraging due to smell or taste changes, although we are also unaware of studies in which this has been evaluated. In contrast to this set of mechanisms by which antibiotics could reduce foraging, the biological spray could have instead increased the attractiveness of the flowers in the field, leading to greater floral visitation. The biological spray active ingredient is a yeast, and honey bees can detect volatiles produced by floral fungi and yeasts (Rering et al., 2018). For example, honey bee visitation to pear flowers increased when the yeast *Metschnikowia* and the bacterium *Acinetobacter nectaris* were sprayed together (Colda et al., 2021). Additionally, though we did not find statistically significant differences in motor activity between control bumble bees and those fed high doses of oxytetracycline for three days, biological-fed bumble bees were more active than antibiotic-fed bumble bees (Fig 5B), indicating no negative effects on locomotion. This finding supports previous evidence that *Bombus terrestris* foraging behavior was unaffected by feeding on *A. pullulans* (Iqbal et al., 2022). Overall, our behavioral findings agree with previous lab experiments demonstrating that exposure to antibiotics can alter bee foraging behavior, but further studies are needed to understand these mechanisms (but see Ortiz-Alvarado et al., 2022).

Impacts on bee foraging, like those seen in our experiments, can have cascading consequences for colony development. The effects may not manifest in the systems where the impact occurred (in this case, pears), but may have economic consequences for subsequent crops using those bees and for beekeepers. Past work has demonstrated that decreases in foraging efficiency (i.e., lower ratio of pollen collected to foraging time) and reduced floral visitation due to pesticides can negatively impact social bee colonies. Insecticides have been shown to reduce foraging efficiency in bumble bees (Feltham et al., 2014; Gill et al., 2012; Stanley et al., 2016), leading to smaller colonies in some cases (Feltham et al., 2014). Fungicides have also been shown to reduce the number of foraging bouts, having negative impacts on honey bee pollen storage (Prado et al., 2019). Therefore, a future direction of this work would be to examine the effects of fire blight materials on colony health.

Honey bee foraging declines in the field did not translate to pollination and fruit quality reductions in our study, suggesting that using these fire blight sprays may not have direct economic consequences for pear growers. First, we did not see differences among the stigmatic pollen deposition in unbagged branches between the two fire blight treatments (Figure 2A). The lack of differences in pollen deposition among treatments was surprising because we observed fewer flower visits by bees in the antibiotic-sprayed sites. This result could be due, in part, to the extended length of the bee visits to flowers at antibiotic-sprayed sites (Figure 1B), which could have increased the transfer of pollen from anthers to stigmas despite fewer visits. Indeed, the average number of pollen grains present in stigmas from both treatments is in line with reports of previous pear studies (Bieniasz et al., 2017) and above the minimum number needed to fertilize all five floral carpels. It is important to remember that, to study the realistic potential for self-pollination in the field, we did not remove anthers were not removed prior to flower bagging. Second, we did not see any treatment effects when we analyzed the proportion of flowers with ovule expansion on those same branches (Figure 2B). Approximately 40% of the flowers bagged until petal fall exhibited ovule expansion (Figure S5), which indicates a high degree of parthenocarpy or/and selfing (Nishitani et al., 2012; Quinet and Jacquemart, 2015). Given that we did not record the tree cultivar, it is impossible to know whether fruits in bagged branches developed due to parthenocarpy or to self-fertilization (which is known to happen in Bartlett pears, Burts and Kelly, 1960). Third, we detected a statistically significant (p<0.001), but modest effect of insect activity on ovule expansion (∼24% increase in open relative to bagged branches). This is likely due to the unusually cold Spring of 2022 in central Washington State, which impacted bee foraging activity (Sallato and Whiting, 2022) and could have had negative impacts on pollen tube germination (Bieniasz et al., 2017; Claessen et al., 2019; Kuroki et al., 2017).

Given the pollination results, unsurprisingly, the fire blight treatments did not affect the measured agronomic outcomes. We harvested and counted the number of mature seeds in D’Anjou pears, which require insect cross-pollination. There were no differences in the number of seeds among unsprayed control, antibiotic, and biological-sprayed sites (Figure 3A). The seed set results, however, could be biased, as trees could have abscised poorly fertilized fruits prior to fruit harvest (Webster, 2002), and often orchardists hand-thin small fruit off the tree. The fact that most fruits did not reach their theoretical maximum number of seeds (flowers have five carpels with two ovules each) indicates some level of pollination limitation, although it would be difficult to know if this was due to lack of cross-pollination or due to the cold spring. We also measured fruit diameter, length, and weight in fruits. Similarly to Callan and Lombard (1978), we found that pear fruit diameter increased with the presence of seeds (Figure S6). This is perhaps due to the release of auxins which in turn cause the cells in pomme fruits to grow larger (Devoghalaere et al., 2012). An increase in fruit size, even if modest, could potentially lead to a greater yield per acre, bolstering the profitability of pear operations.

Future field studies should address aspects not considered here. First, we are likely reporting on the lower range of impacts of fire blight treatment on field honey bee behavior. In our experiments, growers applied the antibiotic and the biological antagonist only once due to low disease pressure during an abnormally cold spring. However, management programs in warmer spring seasons in the Pacific Northwest, and typically in the eastern US regions (Martinez, 2017; Walgenbach et al., 2021), include repeated applications of multiple types of antibiotics. Assessing the measured outcomes in years with warmer spring conditions would be helpful, as disease incidence might require more sprays and bee activity might be higher. Second, based on the foraging changes we detected, it is essential to further assess any long-term colony-level impacts, as the changes could affect bee larval development (Duan et al., 2021) and thus overall colony growth, vigor, and fecundity. Third, future studies should account for the differences in pear pollination across varieties (Nishitani et al., 2012). Fourth, studies should consider aspects of tree phenology to assess agronomic outcomes. For example, we only assessed seed set and fruit quality at harvest time, but it is necessary to measure fruit development at different points of the growing season such as pre and post-fruit drop (Callan and Lombard, 1978). Finally, it is crucial to examine how changes in bee behavior could impact other pollinator-dependent crops, such as apples (Garratt et al., 2014), which receive more antibiotic sprays than pears (Figure S1A).

## 5. Conclusion

Bacterial disease management is key to pome fruit production, yet it carries unknown consequences for beneficial insects and pollination services. While there are few laboratory studies on the impact of bactericide treatments on pollinators, all indicate that antibiotics could impact bee foraging (Avila et al., 2022; Ortiz-Alvarado et al., 2020). Our field-based results are consistent with these laboratory studies. Honey bees visited fewer flowers after the orchards were sprayed with oxytetracycline, while the opposite was true for the biological antagonist *Aureobasidium pullulans*. Additionally, given the complexities involved, very few studies assessing the impacts of agricultural inputs more broadly have tried correlating behavioral changes to pollination outcomes (but see, e.g., Stanley et al., 2015, focused on neonicotinoid insecticides). Here we report that the behavioral changes we detected did not scale up to create detectable impacts on pear pollination and fruit set. Still, we assessed an atypically field-realistic agricultural exposure (a single spray) and worked in pears, many cultivars of which can self-fertilize.

More research is needed to fully assess the impact of field-realistic fire blight treatment regime on colony health, bee fitness, and crop production. For example, we need to determine the temporal extent of the impacts on honey bee foraging. The changes could, for example, affect multiple bee generations. Additionally, honey bee functioning, and health could be compromised by the depletion of core gut symbionts (Raymann and Moran, 2018). Such field-realistic research could inform management recommendations to beekeepers, growers, and policymakers. This is especially important in light of the continued use of antibiotics in pome fruits in the USA (Figure S1) and the expansion of fire blight to other regions in the world (Zhao et al., 2019).

## Data availability

data and code are deposited at Figshare and will be available upon review.

## Acknowledgements

we would like to thank all the anonymous growers who participated in the study; members of the Nottingham lab who harvested and processed the pear fruits; Dr. Lee Kalcsits for fruit quality measurements; Joselyne Chavez and James Allen for support with the research logistics; Sherry Tsiu, Rohan Singh, and Alexa McGrath for helping with some of the stigmatic pollen deposition assessment; and Jeeya Sharma for the graphical abstract.

