## Supplementary Figures and Tables for "Effects of a field-sprayed antibiotic on bee foraging behavior and pollination in pear orchards"

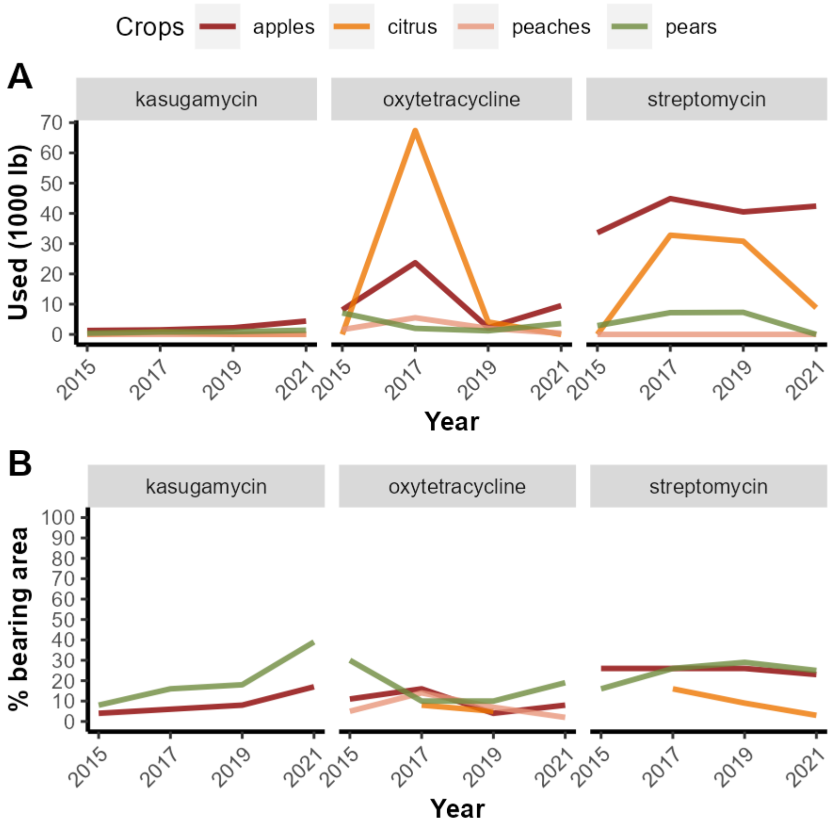

**Figure S1. Use of crop antibiotics in the United States since 2015.** A) Antibiotic usage in the US by crop; B) Area under cultivation sprayed with antibiotics. Data mined from the National Agricultural and Statistics Service of the United States Department of Agriculture (National Agricultural Statistics Service, 2022). Antibiotic applications were used up until 2014 in organic orchards (Godoy, 2013), hence here we present data from 2015 onwards.

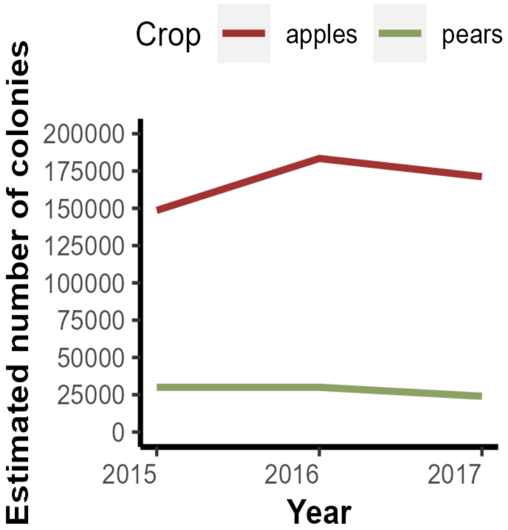

**Figure S2**. Estimated number of honey bees colonies placed in apple and pear orchards from 2015 to 2017 (most recent data reported). Data mined from the National Agricultural and Statistics Service of the United States Department of Agriculture (National Agricultural Statistics Service, 2022).

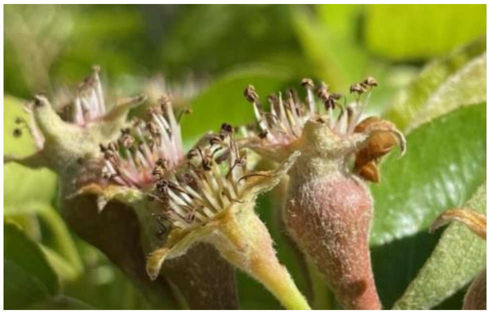

**Figure S3. Picture of flowers with and without ovule expansion.** Pictures taken at petal fall.

Godoy, M., 2013. A Battle Over Antibiotics In Organic Apple And Pear Farming [WWW Document]. Salt - Natl. Public Radio. URL https://www.npr.org/sections/thesalt/2013/04/08/176606069/surprise-organic-apples-and-pears-aren-t-free-of-antibiotics (accessed 2.13.23).

National Agricultural Statistics Service, 2022. USDA/NASS QuickStats Ad-hoc Query Tool [WWW Document]. Agric. Chem. Use Progr. URL https://quickstats.nass.usda.gov/ (accessed 2.13.23).

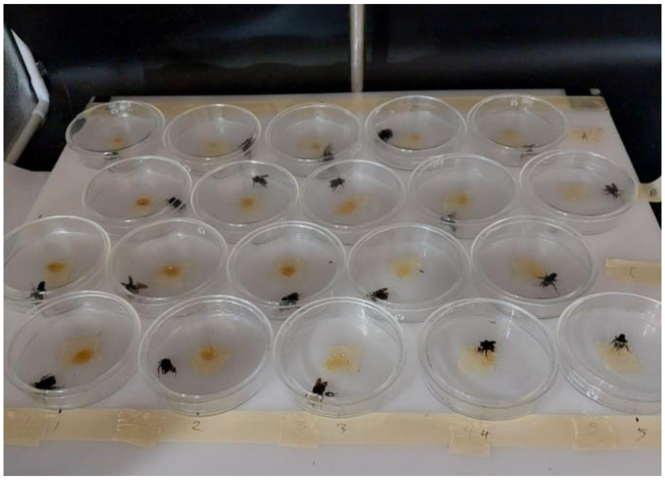

**Figure S4. Artificial foraging arenas monitored for 45 minutes.** Bees were placed within 10 mm Petri dishes with a droplet of honey in the middle (reward). We recorded bee behavior for 45 minutes through an EthoVision XT (Noldus Information Technology, Leesburg, VA, USA).

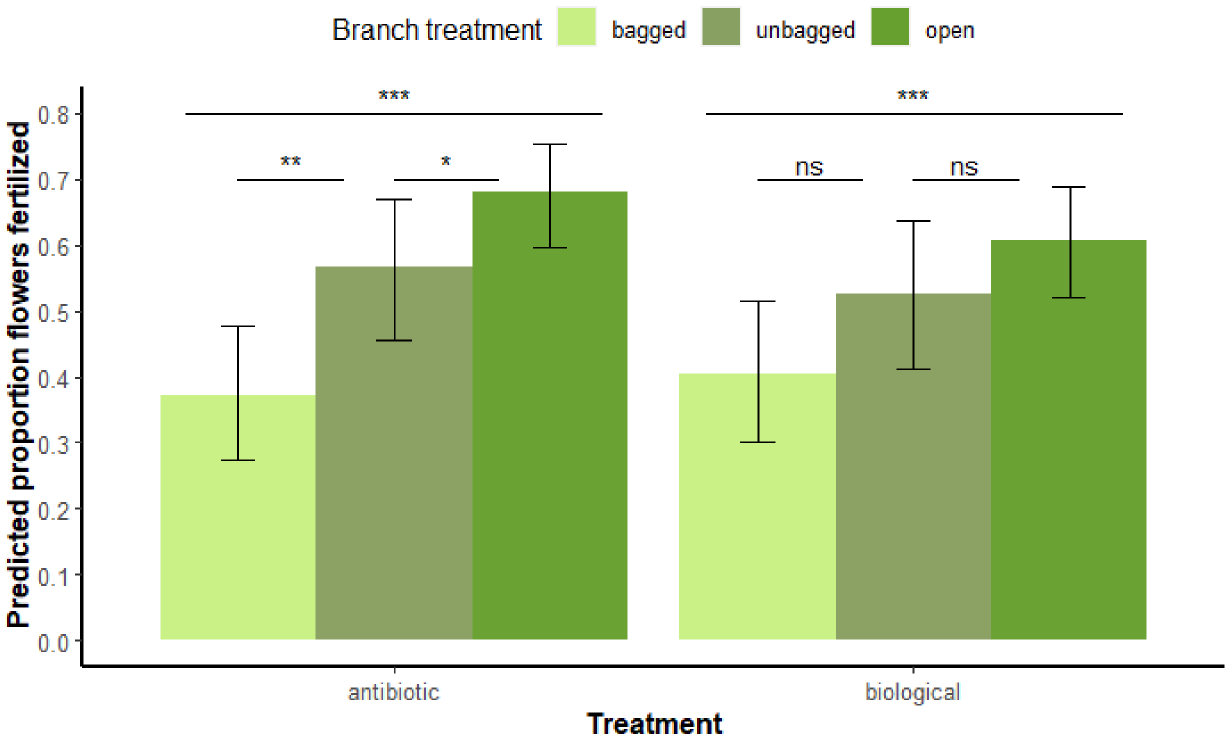

**Figure S5. Pears have a high degree of parthenocarpy and/or self-pollination.** Bagged are branches in muslin bags until petal fall, unbagged are branches where bags were removed right after antibiotic sprays until petal fall, open branches remained open until petal fall. Bars indicate means plus 95% CI. Asterisks indicate statistically significant differences against control, with * = $p < 0.05$; ** = $p < 0.01$; *** = $p < 0.001$

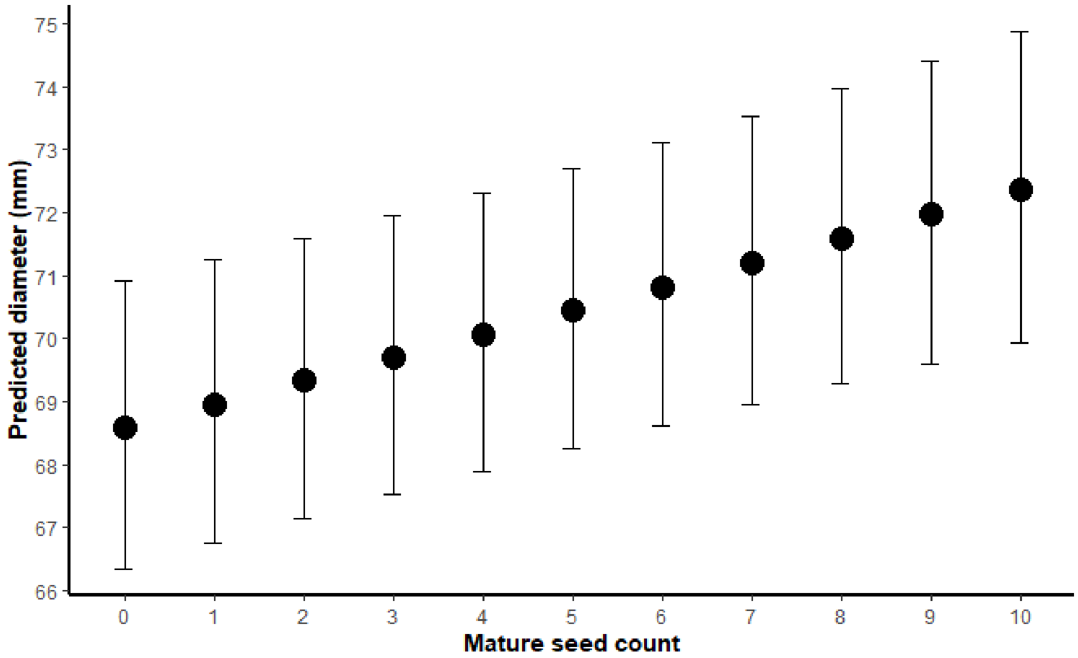

**Figure S6. Predicted increase in D’Anjou pear diameter with increasing mature seed number.**

### Supplementary Tables

Table S1. GLMM model for number of flowers visited per field honey bee forager.

| Predictors | Incidence Rate Ratios (Odds Ratio) | CI | *P* |
|---|---|---|---|
| (Intercept) | 0.14 | 0.09 – 0.20 | <0.001 |
| *Fixed effects* | | | |
| Treatment: antibiotic | 1.47 | 1.08 – 2.01 | 0.016 |
| Timing: post | 1.24 | 0.99 – 1.55 | 0.057 |
| Timing: post2 | 1.48 | 0.98 – 2.24 | 0.064 |
| Flower density | 1 | 1.00 – 1.00 | 0.05 |
| Honey bee abundance | 1.01 | 1.00 – 1.02 | 0.141 |
| Treatment: antibiotic x Timing: post | 0.69 | 0.57 – 0.84 | <0.001 |
| Treatment: antibiotic x Timing: post2 | 0.45 | 0.28 – 0.72 | 0.001 |
| *Random Effects* | | | |
| σ2 | 2.87 | | |
| τ00 Sites | 0.03 | | |
| ICC | 0.01 | | |
| Sites | 10 | | |
| Observations (bees that visited flowers) | 443 | | |
| *R2 marginal* | 0.14 | | |
| *R2 conditional* | 0.29 | | |

Table S2. Wald $X^2$ for number of flowers visited per field honey bee forager.

| Predictor | Chi Square | DF | *P* |
|---|---|---|---|
| Treatment | 5.844 | 1 | 0.01563 |
| Timing | 4.092 | 2 | 0.129237 |
| Flower density | 3.839 | 1 | 0.050076 |
| Honey bee abundance | 2.166 | 1 | 0.141109 |
| Treatment x Timing | 16.796 | 2 | 0.000225 |

Table S3. GLMM model for length of visit of field honey bee forager per flower.

| Predictors | Estimate | CI | *P* |
|---|---|---|---|
| (Intercept) | 2.07 | 1.68 – 2.46 | <0.001 |
| *Fixed effects* | | | |
| Treatment: antibiotic | -0.32 | -0.62 – -0.02 | 0.037 |
| Timing: post | 0.01 | -0.26 – 0.29 | 0.916 |
| Timing: post2 | -0.1 | -0.43 – 0.24 | 0.577 |
| Flower density | 0 | -0.00 – 0.00 | 0.285 |
| Treatment: antibiotic x Timing: post | 0.36 | 0.07 – 0.66 | 0.016 |
| Treatment: antibiotic x Timing: post2 | 0.67 | 0.19 – 1.16 | 0.007 |
| *Random Effects* | | | |
| σ2 | 0.42 | | |
| τ00 Sites | 0.02 | | |
| ICC | 0.04 | | |
| Sites | 10 | | |
| Observations (bees that visited flowers) | 443 | | |
| *R2 marginal* | 0.05 | | |
| *R2 conditional* | 0.09 | | |

Table S4. Wald $X^2$ for length of visit of field honey bee forager per flower.

| Predictor | Chi Square | DF | *P* |
|---|---|---|---|
| Treatment | 4.349 | 1 | 0.0370369 |
| Timing | 0.876 | 2 | 0.6454392 |
| Flower density | 1.141 | 1 | 0.2854467 |
| Treatment x Timing | 8.911 | 2 | 0.0116149 |

Table S5. GLMM model for proportion number of pollen grains deposited per pear flower stigma.

| Predictors | Incidence Rate Ratios (Odds Ratio) | CI | *P* |
|---|---|---|---|
| (Intercept) | 69.19 | 44.43 – 107.77 | <0.001 |
| *Fixed effects* | | | |
| Treatment: antibiotic | 0.94 | 0.50 – 1.76 | 0.8490 |
| *Random Effects* | | | |
| σ2 | 0.23 | | |
| τ00 Unique Trees:Site | 0.06 | | |
| τ00 Site | 0.13 | | |
| ICC | 0.47 | | |
| Unique branches | 59.00 | | |
| Sites | 6 | | |
| Observations (Flowers) | 117 | | |
| *R2 marginal* | 0.00 | | |
| *R2 conditional* | 0.47 | | |
| *R2 marginal* | 0.002 | | |
| *R2 conditional* | 0.49 | | |

Table S6. Wald $X^2$ for proportion number of pollen grains deposited per pear flower stigma.

| Predictor | Chi Square | DF | *P* |
|---|---|---|---|
| Treatment | 0.034 | 1 | 0.848700 |

Table S7. GLMM model for proportion of pear flowers with ovule expansion per branch.

| **Predictors** | **Incidence Rate Ratios (Odds Ratio)** | **CI** | ***P*** |
|---|---|---|---|
| (Intercept) | 0.84 | 0.54 – 1.31 | 0.4350 |
| *Fixed effects* | | | |
| Treatment: antibiotic | 0.96 | 0.54 – 1.71 | 0.8990 |
| Branch: open | 1.94 | 1.43 – 2.62 | <0.001 |
| Branch: unbagged | 1.57 | 1.07 – 2.31 | 0.0230 |
| Clusters Total Count | 0.92 | 0.88 – 0.96 | <0.001 |
| Antibiotic x open | 1.45 | 0.96 – 2.18 | 0.0780 |
| Antibiotic x unbagged | 1.14 | 0.67 – 1.93 | 0.6340 |
| | 140.53 | 58.10 – 411.94 | |
| *Zero-Inflated Model* | | | |
| (Intercept) | 0.03 | 0.02 – 0.05 | <0.001 |
| *Random Effects* | | | |
| σ2 | -0.02 | | |
| τ00 Unique Trees:Site | 0.1 | | |
| τ00 Site | 0.13 | | |
| ICC | 1.07 | | |
| Unique Trees | 367 | | |
| Sites | 10 | | |
| Observations (Flowers) | 510 | | |

Table S8. Wald $X^2$ for proportion of pear flowers with ovule expansion per branch.

| **Predictor** | **Chi Square** | **DF** | ***P*** |
|---|---|---|---|
| Treatment | 0.859 | 1 | 0.353928 |
| Branch Treatment | 67.209 | 2 | 0.000000 |
| Number of clusters/branch | 13.014 | 1 | 0.000309 |
| Treatment x Branch Treatment | 3.784 | 2 | 0.150796 |

Table S9. GLMM model number of seeds in mature fruits of D'Anjou pear cultivar.

| **Predictors** | **Incidence Rate Ratios (Odds Ratio)** | **CI** | ***P*** |
|---|---|---|---|
| (Intercept) | 4.43 | 2.85 – 6.86 | <0.001 |
| *Fixed effects* | | | |
| Treatment: antibiotic | 0.93 | 0.53 – 1.65 | 0.8150 |
| Treatment: biological | 0.97 | 0.46 – 2.01 | 0.9280 |
| Cultivar:Red_Anjou | 0.75 | 0.41 – 1.37 | 0.3500 |
| *Random Effects* | | | |
| σ2 | 0.25 | | |
| *τ00 Sites* | 0.14 | | |
| ICC | 0.36 | | |
| Sites | 13 | | |
| Observations (Fruits) | 648 | | |
| *R2 marginal* | 0.05 | | |
| *R2 conditional* | 0.41 | | |

Table S10. Wald $X^2$ for number of seeds in mature fruits of D'Anjou pear cultivar.

| **Predictor** | **Chi Square** | **DF** | ***P*** |
|---|---|---|---|
| Treatment | 0.059 | 2 | 0.971079 |
| Cultivar | 0.874 | 1 | 0.349909 |

Table S11. GLMM model for fruit diameter in mature fruits of D’Anjou pear cultivar.

| Predictors | Estimate (log) | CI | *P* |
|---|---|---|---|
| (Intercept) | 4.24 | 4.20 – 4.28 | <0.001 |
| *Fixed effects* | -0.01 | -0.06 – 0.04 | 0.7390 |
| Treatment: antibiotic | -0.02 | -0.08 – 0.04 | 0.5560 |
| Treatment: biological | 0.02 | -0.04 – 0.07 | 0.5620 |
| Cultivar:Red_Anjou | 0.01 | 0.00 – 0.01 | <0.001 |
| *Random Effects* | | | |
| σ2 | 0 | | |
| τ00 Sites | 0 | | |
| ICC | 0.2 | | |
| Sites | 13 | | |
| Observations | 648 | | |
| *R2 marginal* | 0.04 | | |
| *R2 conditional* | 0.23 | | |

Table S12. Wald $X^2$ for fruit diameter in mature fruits of D’Anjou pear cultivar.

| Predictor | Chi Square | DF | *P* |
|---|---|---|---|
| Treatment | 0.347 | 2 | 0.840685 |
| Cultivar | 0.336 | 1 | 0.562182 |
| Mature seed count | 19.277 | 1 | 0.000011 |

Table S13. GLMM model for fruit length in mature fruits of D’Anjou pear cultivar.

| Predictors | Estimates (sqrt) | CI | *P* |
|---|---|---|---|
| (Intercept) | 9.97 | 9.76 – 10.18 | <0.001 |
| *Fixed effects* | | | |
| Treatment: antibiotic | 0.06 | -0.21 – 0.33 | 0.669 |
| Treatment: biological | -0.17 | -0.52 – 0.18 | 0.348 |
| Cultivar: red D'Anjou | 0.15 | -0.14 – 0.44 | 0.303 |
| *Random Effects* | | | |
| σ2 | 0.24 | | |
| *τ00 Sites* | 0.03 | | |
| ICC | 0.11 | | |
| Sites | 13 | | |
| Observations (Fruits) | 598 | | |
| *R2 marginal* | 0.02 | | |
| *R2 conditional* | 0.13 | | |

Table S14. Wald $X^2$ for fruit length in mature fruits of D'Anjou pear cultivar.

| Predictor | Chi Square | DF | *P* |
|---|---|---|---|
| Treatment | 2.398 | 2 | 0.301441 |
| Cultivar | 1.060 | 1 | 0.303322 |

Table S15. GLMM model for fruit weight in mature fruits of D'Anjou pear cultivar.

| Predictors | Estimates (sqrt) | CI | *P* |
|---|---|---|---|
| (Intercept) | 13.67 | 12.95 – 14.38 | <0.001 |
| *Fixed effects* | -0.37 | -1.56 – 0.82 | 0.54 |
| Treatment: antibiotic | -0.03 | -1.15 – 1.10 | 0.963 |
| Treatment: biological | 0.27 | -0.92 – 1.46 | 0.654 |
| Cultivar: red D'Anjou | | | |
| *Random Effects* | | | |
| σ2 | 1.87 | | |
| *τ00 Sites* | 0.55 | | |
| ICC | 0.23 | | |
| Sites | 13 | | |
| Observations (Fruits) | 598 | | |
| *R2 marginal* | 0.01 | | |
| *R2 conditional* | 0.23 | | |

Table S16. Wald $X^2$ for fruit weight in mature fruits of D'Anjou pear cultivar (n = 648 pear fruits).

| Predictor | Chi Square | DF | *P* |
|---|---|---|---|
| Treatment | 0.383 | 2 | 0.825550 |
| Cultivar | 0.201 | 1 | 0.654162 |

Table S17. GLMM model for number of approaches to reward by individuals of *Bombus vosnesenskii* over 45 minutes trial.

| Predictors | Incidence Rate Ratios (Odds Ratio) | CI | *P* |
|---|---|---|---|
| (Intercept) - control sucrose | 76.4 | 51.95 – 112.36 | <0.001 |
| *Fixed Effects* | | | |
| Treatment: biological full rate | 0.58 | 0.37 – 0.91 | 0.019 |
| Treatment: full antibiotic rate | 0.63 | 0.40 – 0.99 | 0.044 |
| Treatment: half antibiotic rate | 0.49 | 0.31 – 0.78 | 0.002 |
| *Random Effects* | | | |
| $\sigma^2$ | 0.43 | | |
| $\tau_{00}$ Colony:Quad | 0 | | |
| $\tau_{00}$ Quad | 0.05 | | |
| Colonies | 16 | | |
| Quads | 4 | | |
| N Bees | 80 | | |
| *R2 marginal* | 0.13 | | |
| *R2 conditional* | 0.22 | | |

Table S18. Wald $X^2$ for number of approaches to reward by individuals of *Bombus vosnesenskii* over 45 minutes trial.

| Predictor | Chi Square | DF | *P* |
|---|---|---|---|
| Treatment | 10.275 | 3 | 0.016360 |

Table S19. GLMM model for amount of spent moving by individuals of *Bombus vosnesenskii* over 45 minutes trial.

| **Predictors** | **Incidence Rate Ratios (Odds Ratio)** | **CI** | ***P*** |
|---|---|---|---|
| (Intercept) - control sucrose | 707.24 | 323.63 – 1090.85 | <0.001 |
| *Fixed Effects* | 163.11 | -241.21 – 567.43 | 0.429 |
| Treatment: biological full rate | -122.43 | -526.75 – 281.89 | 0.553 |
| Treatment: full antibiotic rate | -223.29 | -627.61 – 181.03 | 0.279 |
| Treatment: half antibiotic rate | | | |
| *Random Effects* | | | |
| σ2 | 251233.67 | | |
| τ00 Colony:Quad | 34864 | | |
| τ00 Quad | 68119.52 | | |
| ICC | 0.29 | | |
| Colony | 16 | | |
| Quad | 4 | | |
| Bees | 80 | | |
| *R2 marginal* | 0.06 | | |
| *R2 conditional* | 0.33 | | |

Table S20. Wald Wald $X^2$ for time spent in motion by *Bombus vosnesenskii* over 45 minutes trial.

| **Predictor** | **Chi Square** | **DF** | ***P*** |
|---|---|---|---|
| Treatment | 3.906 | 3 | 0.271800 |

Table S21. Summary of data collected per experimental site (orchard).

| **Site Code** | **Treatment** | **Selection & bagging of branches** | **Pre spray honey bee sampling & bee abundance** | **Spray date** | **Post honey bee sampling & bee abundance** | **Post spray bag removal** | **Post spray stigmas collection** | **Post2 honey bee sampling & bee abundance** | **Ovule expansion assessment** | **Pear harvest** |
|---|---|---|---|---|---|---|---|---|---|---|
| 1 | Antibiotic | 4/6/2022 | 4/19/2022 | 4/22/2022 | 4/23/2022 | 4/23/2022 | 4/26/2022 | 4/26/2022- only 2 bee | 5/10/2022 to 5/11/22 | 9/13/2022 to 9/27/2022 |
| 2 | Antibiotic | 4/11/2022 | 4/22/2022 | 4/26/2022 | 4/27/2022 | 4/27/2022 | 5/3/2022 | 5/3/2022 | 5/10/2022 to 5/11/22 | 9/13/2022 to 9/27/2022 |
| 3 | Antibiotic | 4/13/2022 | 4/26/2022 | 5/3/2022 | 5/4/2022 | 5/4/2022 | . | . | 5/10/2022 to 5/11/22 | 9/13/2022 to 9/27/2022 |
| 4 | Antibiotic | 4/11/2022 | 4/25/2022 | 5/3/2022 | 5/4/2022 | 5/4/2022 | . | . | 5/10/2022 to 5/11/22 | 9/13/2022 to 9/27/2022 |
| 5 | Antibiotic | 4/8/2022 | no bees - cold | 4/19/2022 | 4/20/2022 | 4/20/2022 | 4/25/2022 | 4/25/2022 | 5/10/2022 to 5/11/22 | 9/13/2022 to 9/27/2022 |
| 1 | Biological | 4/7/2022 | 4/26/2022 | 4/27/2022 | 4/28/2022 | 4/27/2022 | . | . | 5/10/2022 to 5/11/22 | 9/13/2022 to 9/27/2022 |
| 2 | Biological | 4/6/2022 | 4/19/2022 | 4/19/2022 | 4/20/2022 | 4/20/2022 | 4/25/2022 | 4/25/2022 | 5/10/2022 to 5/11/22 | 9/13/2022 to 9/27/2022 |
| 3 | Biological | 4/12/2022 | no bees yet | 4/22/2022 | 4/23/2022 | 4/23/2022 | 4/27/2022 | 4/27/2022 | 5/10/2022 to 5/11/22 | 9/13/2022 to 9/27/2022 |
| 4 | Biological | 4/12/2022 | no bees yet | 4/22/2022 | 4/23/2022 | 4/23/2022 | 4/27/2022 | 4/27/2022 | 5/10/2022 to 5/11/22 | 9/13/2022 to 9/27/2022 |
| 5 | Biological | 4/14/2022 | 4/28/2022 | 4/29/2022 | 4/29/2022 | . | . | . | 5/10/2022 to 5/11/22 | 9/13/2022 to 9/27/2022 |
| 6 | Biological | . | . | 4/22/2022 | . | . | . | . | . | 9/13/2022 to 9/27/2022 |

Table S22. Pollination raw data summarized by branch treatment.

| **Branch Treatment** | **Mean proportion** | **Standard Deviation** |
|---|---|---|
| Bagged | 0.366 | 0.264 |
| Open | 0.603 | 0.258 |
| Unbagged | 0.498 | 0.282 |
